# DNA synthesis inside the hepatitis B virus creates a high-energy spool

**DOI:** 10.64898/2026.01.22.701187

**Authors:** Nora Gibes, Katie Culhane, Haitao Liu, Ji Xi, Karolyn Pionek, Smita Nair, Daniel D Loeb, Jianming Hu, Adam Zlotnick, Joseph Che-Yen Wang

## Abstract

The Hepatitis B Virus (HBV) capsid assembles around an RNA pregenome which is reverse-transcribed into double-stranded DNA. It remains unclear how this DNA, which is stiff, bulky, and has significant negative charge, is accommodated within the capsid. The organization of the genome will answer this and lend insight into the viral reverse transcription reaction. We determined the structures of mature HBV and Duck HBV, finding that the DNA forms a spool that is coaxial with the capsid’s fivefold symmetry and interacts with charges on the capsid interior. We calculate that the energy of DNA bending would create a metastable capsid that just awaits a trigger for DNA release. Thus, we see how nature has created a fragile energy minimum that enables genome uncoating.

## Introduction

Hepatitis B Virus (HBV) has infected about 2 billion people, leaving more than 250 million individuals with chronic HBV and causing about one million deaths annually^1^. Like other members of the *Hepadnaviradae*, HBV packages an mRNA pre-genome (pgRNA) that is the template for production of a mature relaxed circular (rcDNA) genome. Reverse transcription of the linear single-stranded pgRNA to produce the double-stranded mature rcDNA genome is a multi-step process that occurs within the viral capsid.^2^ It considerably alters the physical-chemical nature of the packaged nucleic acid. While the net charge of pgRNA is approximately neutralized by positive charges on the inner surface of the capsid, rcDNA production roughly doubles the nucleic acid charge. Also, as a double-stranded DNA (dsDNA) molecule with short single-stranded gaps, rcDNA is a stiff polymer with a much longer persistence length compared to pgRNA (50 nm vs. 1-2 nm)^3–5^. As a result, fitting rcDNA into the HBV capsid, with its internal diameter of only 25 nm, creates a high-energy state that may contribute to the observed fragility of the mature nucleocapsid^6^. Disassembly at the right time and place is essential for virus replication, and this stress from rcDNA may be an important factor that modulates capsid stability.

The problem of packaging dsDNA within a viral capsid is not unique to *Hepadnaviradae*. Many previous studies have explored how viral DNA becomes organized in the face of the DNA-DNA electrostatic repulsion and bending forces that resist its compaction^7,8^. Potential organizations include a multi-layered coaxial spool, a concentric (but not coaxial) spool, a toroid, and globally disordered loops with local strand alignment.^7,9^ The coaxial spool has been seen in cryogenic electron microscopy (cryo-EM) structures of herpes simplex virus (HSV)^10^, dsDNA bacteriophages^11–16^, and the dsRNA phage phi6^17^. In some cases, symmetry averaging can lead to genomes that appear more ordered than they actually are^9^. Molecular modeling studies^7,9,18^, focused on the phage system, generally favor more disordered arrangements, and these models are also supported by experimental results^19–23^. However, generalizing these studies to the hepadnaviruses is difficult. Most dsDNA viruses and phages use a molecular motor to pump DNA through a portal and into a large-diameter chamber with a relatively neutral charge. In comparison, the hepadnaviral genome is synthesized directly inside a much smaller capsid, with an inner surface that is highly charged (the HBV capsid protein has a 149-residue assembly domain that forms the contiguous shell and a disordered C-terminal nucleic-acid-binding domain, CTD, with 16 arginine residues located inside the capsid). Understanding the genome structure of *Hepadnaviridae* would therefore expand our knowledge of the biophysics of viral genome packaging. It would also provide a window into HBV genome maturation, a key step in HBV replication for which we have almost no structural knowledge.

We sought to determine the hepadnavirus genome structure by performing cryo-EM on HBV and Duck HBV (DHBV) nucleocapsids purified from hepatoma cell culture. We find that these hepadnavirus capsids contain coaxial layers of ringed density consistent with a spooled genome. The rings appear to be B-form DNA and exhibit a wider strand spacing than seen in bacteriophages. Intriguingly, the spool is oriented with the capsid 5-fold axis. These results have implications for nucleic acid organization in HBV and other spherical viruses, as well as for understanding the reverse transcription process.

## Results

### Genome Structure of Duck HBV

In our effort to determine the genome structure of HBV, we started with the DHBV system, as DHBV’s nucleic acid content is more homogeneous^24^ and produces higher nucleocapsid yields. DHBV and HBV share a similar genome and life cycle^25^. Their nucleocapsids consist of a T=4 icosahedral sphere made of 120 dimers of the capsid protein, with the viral polymerase and genome packaged inside. DHBV has a slightly smaller genome, at 3.0 kB, than HBV at 3.2 kB, but a similar internal capsid diameter, so DHBV rcDNA should have slightly more room within the DHBV capsid.

We expressed and purified DHBV from the stably transfected LMH cell line L59-3. Nucleocapsids purified from the cell lysates were imaged by cryo-EM. Approximately 122,883 capsids were selected after 2D classification (Extended Data Fig. 1), in which broken particles, those overlapping, and those obscured by ice were discarded. The capsid structure was solved to 3.3 Å with icosahedral symmetry imposed (Fig. 1a).

**Fig. 1.**
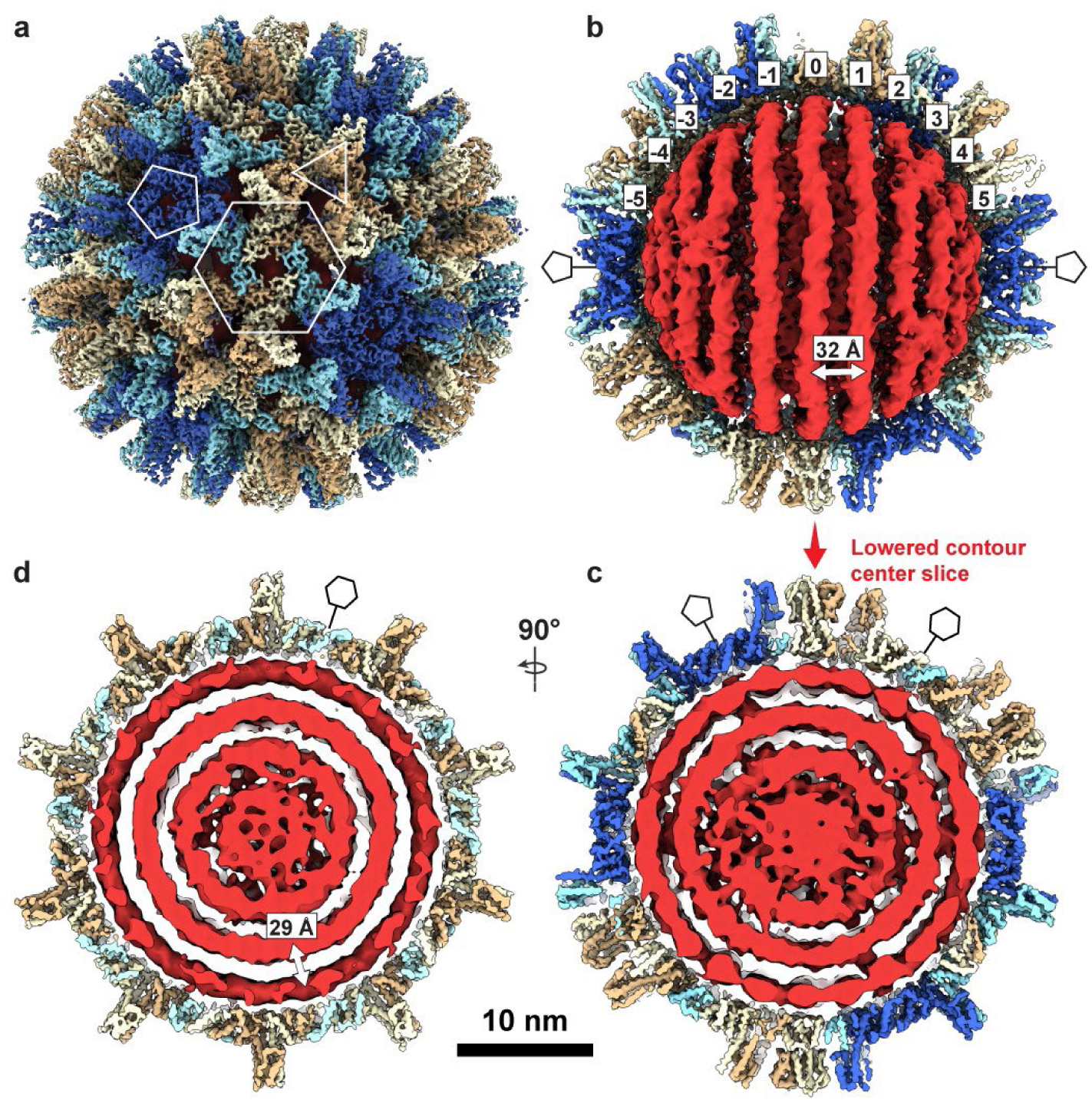
DHBV nucleocapsid structure. **a**, Reconstruction of the T=4 DHBV capsid with icosahedral symmetry imposed. Subunits in the asymmetric unit are colored individually (A, royal blue; B, pale turquoise; C, light cream; D, tan). Quasi-sixfold, threefold, and fivefold overlays (hexagon, triangle, pentagon) indicate positions of the respective symmetry axes. **b**, Genome structure after 3D classification of unsymmetrized subparticles (red). The capsid structure from the corresponding particles is shown cut away (blue and tan).. **c**, A thin, center slice from the structure in a keeping the same alignment (looking down a quasi-six fold), but with lower contouring to show weaker internal density and connecting density between layers. Positions of quasi-sixfold and pentamer pores are indicated along the edge of the capsid. **d**, A thin, center slice from the structure in a looking down a five-fold axis. Positions of quasi-sixfold pores are indicated along the edge of the capsid. Density contouring is the same as in c.

To visualize the non-icosahedral genome structure, we performed multiple rounds of focused 3D classification on the electron density internal to the capsid (Extended Data Fig. 2). This 3D classification was the primary means by which particles were sorted based on the conformational differences in the genome.

We observed several classes (Extended Data Fig. 2) with rings of density consistent with an ordered genome, and these were aligned together to reconstruct the genome (Fig. 1b,c,d red). The resulting structure reveals several rings of density arranged in two spool-like layers, followed by a third, more heterogeneous spherical layer, and an ill-defined central core. DNA density connecting rings and layers has been obscured by positional variance. The concentric rings are coaxially arranged along a 5-fold axis of the icosahedral capsid.

The outermost genome layer is the best-defined, consisting of eleven rings resembling dsDNA strands. We have designated the center ring “0” and numbered the remaining outward as 1-5 (Fig. 1b). Rings “5” and “-5”, directly beneath the capsid 5-fold, could potentially be averaging artifacts, and with a radius of ∼30 Å, they would be around the minimum predicted radius possible for a ring of DNA.^26^ The second layer has at least two apparent coaxial separated rings, with significantly more disorder than those in the first layer (Extended Data Fig. 3a), but the third and the central core lack defined rings and are only visible when rendered at a lowered density contour (Fig. 1c,d, Extended Data Fig. 3a). At low contour one can also observe connections extending from the capsid 5-fold pores through the first and second layers of genome density (Fig. 1c, Extended Data Fig. 3b,c).

### Genome Structure of HBV

We wanted to determine whether this spooled genome is a common feature in HBV as well. DHBV has long been used as a model for understanding reverse transcription in HBV, and a common organization of genomic DNA would further validate that comparison. To obtain mature HBV cores containing rcDNA, we transfected HepG2 cells with a plasmid containing the full HBV genome and purified intracellular nucleocapsids (Extended Data Fig. 4). The preparation contained HBV particles with various genome contents (pgRNA, ssDNA, and dsDNA with a range of completion of the plus strand). Using cryo-EM, we identified a single class that showed the same phenotype of multi-layered concentric ring density observed in DHBV and in tailed dsDNA phages (Extended Data Fig. 4, blue). Without imposing symmetry, a structure based on this class was refined to 3.5Å (Extended Data Table 3). To examine internal features, we used an outside-in strategy: the capsid orientation was fixed, and we first focused on the outer genome layer; once resolved, we proceeded to the second layer (Extended Data Fig. 4, gold) and then to the internal core. Similar to DHBV, the HBV nucleocapsid structure shows a spooled genome aligned with the capsid 5-fold axis, with 9-11 rings in the outer layer (Fig. 2b). The width of these rings is consistent with dsDNA (≈ 20-22Å). The second-layer density is better defined in HBV than in DHBV, and the rings have a width similar to that of the outer layer. In focused classifications, rings from the second layer are seen to pack both behind and between the strands of the outer layers (Extended Data Fig. 5). The resulting coaxial spool is reminiscent of arrangements reported for phage genomes and HSV.^10,13,17^ Three distinct layers are visible in a central slice (Fig. 2d), together with central core density. Again, the third layer is relatively disordered: some regions show thinner density while others show thicker density comparable to the first two layers (Fig. 2d). The central core includes a mass of density that could arise from the viral polymerase (Extended Data Fig. 4). Weak density connecting layers beneath 5-folds is visible at low contour (Fig. 2d), although weaker than in DHBV. One possible source of this attenuation in HBV is particle-to-particle heterogeneity in plus-strand completion and overall genome length. In HBV, the plus strand of rcDNA is typically incomplete and heterogeneous, often only about two-thirds complete, whereas in DHBV, it is nearly complete^27^. Such heterogeneity likely blurs inter-layer density and obscures the apparent polymerase position.

**Fig. 2.**
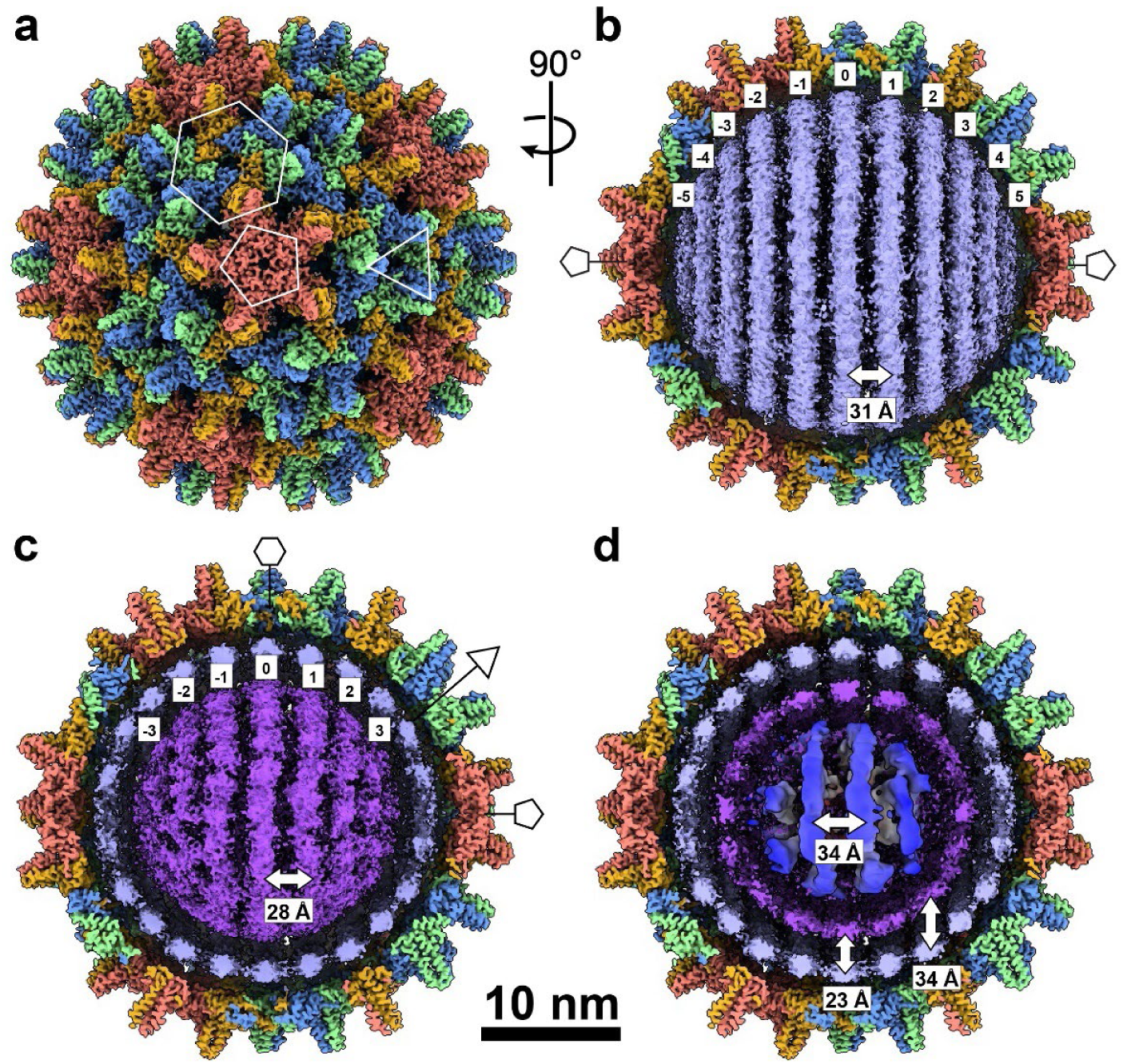
Cryo-EM structure of the mature HBV rcDNA-filled nucleocapsid. **a**, Icosahedrally averaged 3D reconstruction of the HBV T=4 capsid, with core protein subunits in the asymmetric unit colored individually (A, salmon; B, light green; C, cornflower blue; D, goldenrod), and quasi-sixfold, threefold, and fivefold positions marked by a hexagon, triangle, and pentagon, respectively. **b**, Focused classification and 3D refinement of subparticles reveal 11 concentric genome spools (-5 to 5; purple) in the outer DNA layer, shown with the corresponding capsid in cutaway view. The density width is consistent with double-stranded DNA, and the spool at the polar end, nearest the fivefold axis, appears weaker, suggesting increased mobility at this site, potentially modulated by the arginine-rich CTD (Fig. 4). The center-to-center spacing between adjacent spools is ∼31 Å, larger than the ∼25 - 29 Å interaxial spacing reported for dsDNA in tailed bacteriophages (Extended Data Table 2). **c**, Focused classification and 3D refinement at the middle genome layer resolve seven continuous, approximately parallel spools (indexed -3 to 3; purple) with feature widths compatible with double-stranded DNA (Extended Data Fig. 5); the central three are well ordered and show B-form like periodicity, whereas the four spools toward the polar end (two on each side) are more diffuse, likely reflecting interactions with the CTD. The inter-spool spacing is ∼28 Å, indicating tighter packing than in the outer layer. **d**, Central cutaway along the fivefold axis (same contour as **c**), highlighting the innermost core. The nucleic-acid density has a width consistent with a mixture of duplex and single-stranded segments and appears at lower local resolution than the outer layers, indicating greater conformation heterogeneity. A central toroidal density is visible, consistent with the viral reverse transcriptase (Extended Data Fig. 4); interlayer separations vary within this region (for example, ∼23 Å and ∼34 Å; arrows). The capsid layer was reconstructed with icosahedral symmetry (I3); internal genome layers were resolved by focused subparticle analysis asymmetrically (C1).

### Ring density conforms to a B-form DNA structure

The central strands of the outer rings (index number 0 in Fig. 1 and 2) for both DHBV and HBV show repeating surface features consistent with a helical molecule (Fig. 3). The repeat aligns with a B-form double helical nucleic acid model, which has a narrow minor groove and a wide major groove, and a 10 bp repeat that is 33 Å in length (when straight). These repeating features are not consistent with an A-form nucleic acid model, for which the major and minor grooves have similar dimensions and the repeat is 28 Å. The presence of B-form nucleic acid is consistent with dsDNA, and essentially excludes dsRNA or hybrid RNA-DNA as they are expected to exhibit A-form.

**Fig. 3.**
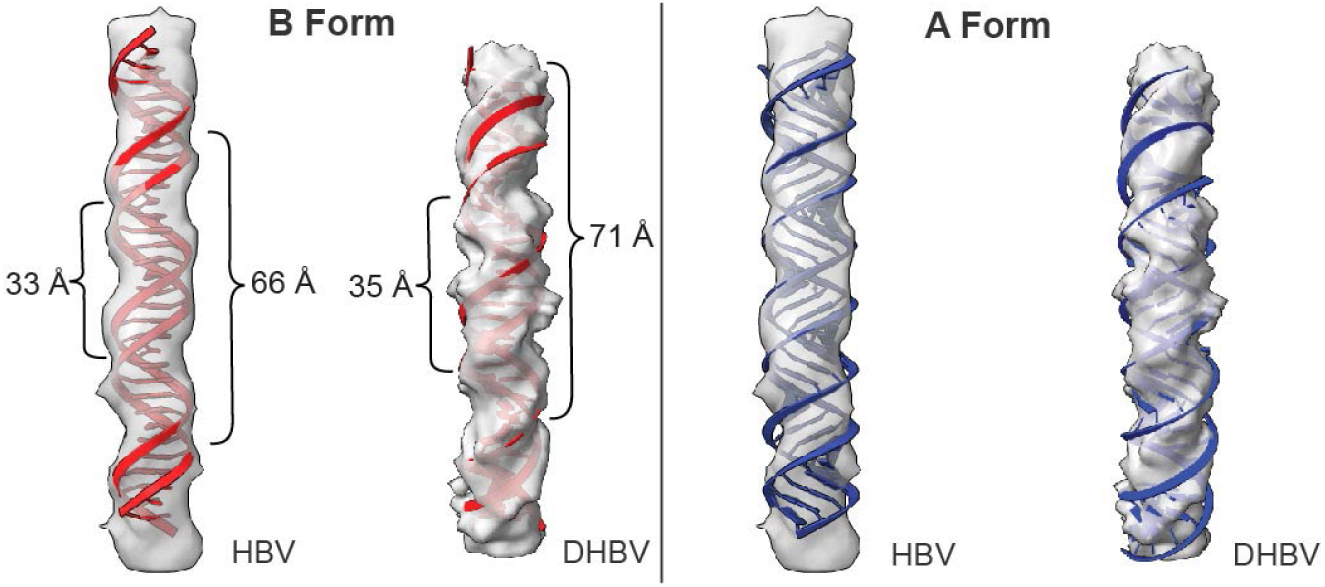
Genome density in HBV and DHBV is compatible with B-form DNA. Representative segments of nucleic-acid density from HBV and DHBV were extracted and fit with idealized duplexes built from a repeating ATCG sequence: B-form models are shown in red (left) and A-form models in blue (right). For HBV, the density periodicity measures ∼33 Å per helical turn and ∼66 Å across two turns, matching the pitch of B-form DNA and mismatching the shorter A-form pitch. For DHBV, the corresponding periodicities are ∼35 Å and ∼71 Å, likewise consistent with B-form. The B-form fits reproduce the groove/crossover positions along the density, whereas A-form models show systematic misalignment and under-pitched helices. Together, these comparisons indicate that the genome segments visualized in both capsids adopt B-form like organization.

## Discussion

We have found that reverse transcription produces a spooled genome for both HBV and DHBV, seen as layers of rings. Density connecting the rings is largely absent; these regions may be mobile or have more random positions, whereas the rings may be fixed in position by interactions with the capsid or represent an averaged position. This organization of the hepadnaviral genome evokes a coiled spring and has implications for DNA packaging in compact spaces, for the process of reverse transcription in hepadnaviruses, the fragility of mature capsids, and the release of packaged DNA.

The outer two layers of the spool have clear rings of dsDNA. A third layer is disordered and may contain some dsDNA. The contour length of the loops in the first two layers could accommodate 2500 bp (HBV) or 2379 bp (DHBV) of DNA (Extended Data Table 1). The fourth layer, the central core, is also disordered; it may contain packaged proteins such as the viral polymerase, host chaperones, helicases, kinases, phosphatases, APOBEC, and other proteins, pro and anti-viral^28–40^. The rings in the outermost layer of the spool are the best resolved and have striations in the density consistent with B-form dsDNA. We cannot exclude the possibility that our reconstruction is contaminated with duplex linear DNA, a minor product resulting from an alternate route in reverse transcription^41^.

The number of rings in the outer layer—9 to 11—aligns with our previous predictions for the HBV genome, which assumed a DNA inter-strand spacing of 26-29 Å^39,42^. While the distance between adjacent layers is roughly 29-30 Å, the ring spacing within the outer layer (inter-strand distance) is 31-32 Å. This is wider interlayer spacing than seen in phages and HSV^10,17,43^, but the inter-strand spacing appears to be similar to HSV (Extended Data Table 2). The wide spacing in the hepadnaviruses may be a result of a lower packing density than many phages (Extended Data Table 2) but could also be impacted by interaction with the arginine-rich CTDs, which are disordered and may extend into the DNA.

While looser spacing would mean reduced electrostatic repulsion from DNA packing, the hepadnaviral DNA strands are still under substantial bending stress due to the small diameter of the capsid lumen (25 nm). This is only half the persistence length of DNA^3^. Stress from genome bending is analogous to a coiled spring and may contribute to the fragility of mature HBV capsids that have been previously observed^6^. We estimate that the outer two layers would generate 217 kcal/mol (HBV) and 197 kcal/mol (DHBV) of bending energy (Extended Data Table 1), or almost 1 kcal/mol/contact, across 240 Cp dimer-dimer interfaces. This would be sufficient to destabilize the capsid, as each interface is only held together by ∼-3 kcal/mol^44^.

We can speculate how reverse transcription in hepadnaviruses leads to the spooled arrangement. We hypothesize that the RNA in the pregenome is positioned around the capsid wall to facilitate template switches during reverse transcription and to avoid tangles (Extended Data Fig. 6 illustrates the reverse transcription process). During negative-strand synthesis, new DNA displaces the RNA. We envision that this requires the polymerase to travel along the RNA strand^45^. Subsequent plus-strand synthesis would produce an initial equatorial ring of dsDNA that would displace ssDNA from the wall (ssDNA displacement is supported by observed APOBEC modifications^39^). The initial rings may slide along the capsid wall to minimize bending energy as more DNA is synthesized. As the first layer becomes fully packed, the DNA rings would settle into the most favorable positions. Based on our structures, we hypothesize that the rings in the outer layer have fixed parallel positions, with mobile connections between rings, rather than being arranged in a continuous climbing spiral.

Perhaps the most intriguing feature of the hepadnaviral DNA spools is that they are coaxial and aligned along the capsid’s fivefold symmetry axis. Evidence of spooled genomes is seen in electron microscopy data for several dsDNA viruses and phages^10–13,15,16,46–50^, and alignment with the fivefold axis is thought to be due to the genome entry portal^51^. However, modeling studies, based on phages with a much larger diameter than that of HBV, including the recent study on HK97^7,9^, suggest instead that many phage genomes are likely to favor a disordered mass of loops, which can appear coaxial and ordered in cryo-EM structures due to reconstruction artifacts from symmetry averaging. When it comes to the hepadnavirus, as the DNA strands in the outer layer of our structures are well ordered enough to resolve B-form striations (Fig. 3, Extended Data Fig. 3b, 4) the organization in at least the outer layer is unlikely to be a product of symmetry averaging.

We suspect two factors may be important in the ordering of the hepadnaviral genome into a coaxial spool. The first is the small size of the hepadnaviral capsid; modeling studies suggest that a smaller capsid interior diameter encourages an ordered spiral genome.^18,52^ The second is interaction with the capsid. In coarse grained models and in phage T7, when the portal complex extends into the center of the capsid, the protrusion appears to guide the genome into a coaxial spool^11,52–54^. However, in bacteriophages (and herpesviruses) the DNA is pumped into a capsid and there is minimal interaction between the negatively charged genome and the capsid wall. In *Hepadnaviradae*, the capsid interior has an icosahedral array of basic amino acids: the arginine-rich Cp CTDs that extend into the capsid lumen. It is known that CTDs can be exposed to the cellular environment by threading through the capsid pores^55^. It is also hypothesized that CTD exposure acts as a signal that impacts capsid trafficking^56^. The positioning of the spool could factor into this process, perhaps leaving CTD more free for exposure and signaling. We see interactions between the CTDs and the packaged dsDNA in our focused reconstructions. In HBV, we observe density connecting the capsid fivefold and sixfold vertices to the first two genome layers (Fig. 4). The fivefold connections are reminiscent of the “stalactite-like” structures found in *in vitro* assembled mutant pgRNA-filled capsids^57^. In our DHBV genome asymmetric reconstruction, this putative CTD density is primarily under the capsid fivefold vertices compared to the sixfold vertices (Extended Data Fig. 3c,d). Potentially, local CTD rearrangements at some sixfold positions are required to accommodate the equatorial DNA ring that runs perpendicular to and underneath the sixfold pores. Consistent with this, the CTD density in HBV is radially concentrated near fivefold axes and more laterally distributed around quasi-sixfold belts (Fig. 4). Interestingly, we previously observed that CTDs under fivefold axes in empty capsids were protected from trypsin digestion^55^. Together, these observations suggest that DNA-CTD interactions may help orient and tether the rings of the outer DNA layer perpendicular to a fivefold symmetry axis and contribute to overall genome organization.

**Fig. 4.**
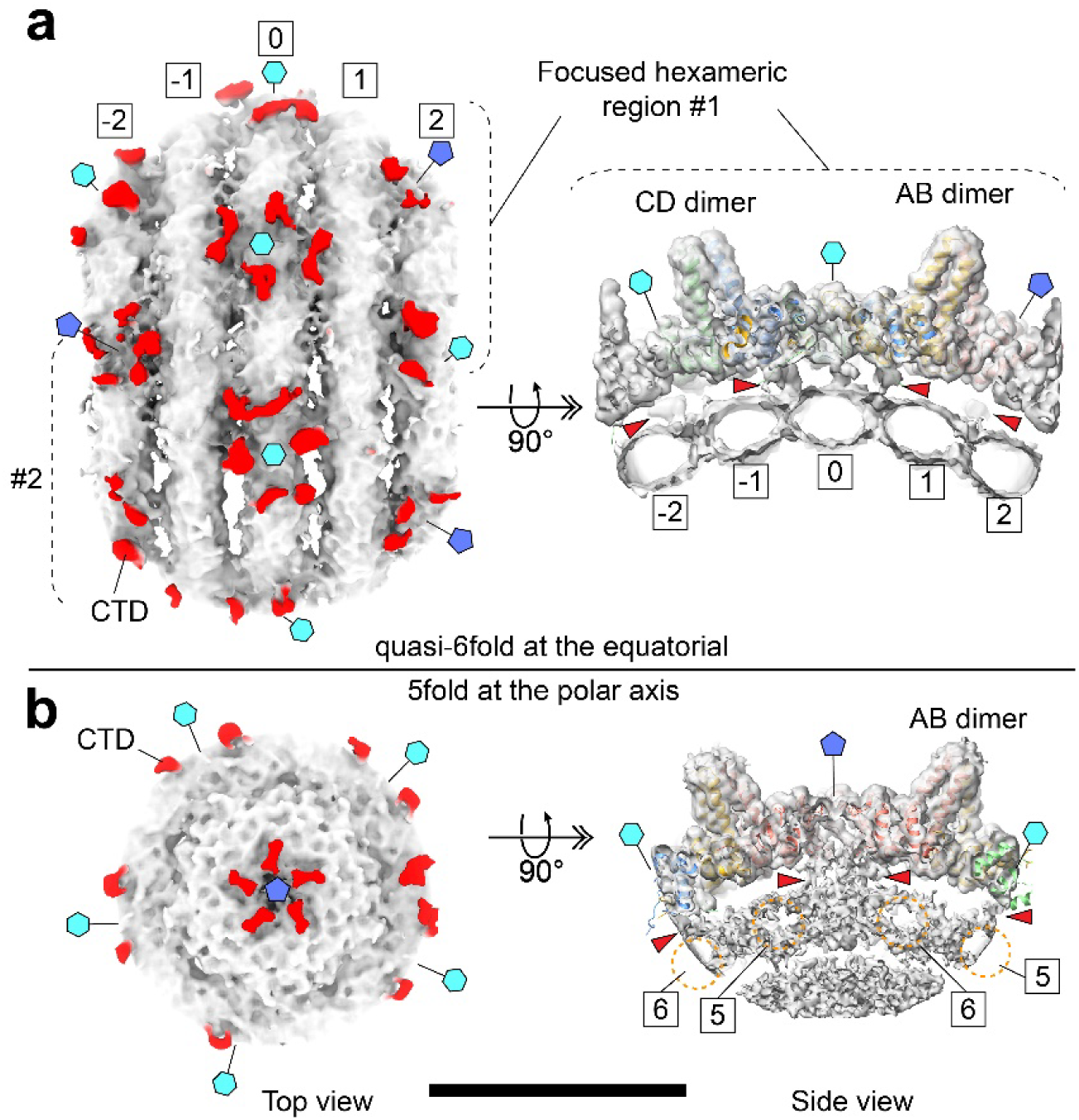
CTD contacts with the outer genome spool at quasi-sixfold and fivefold sites. **a,** Focused subparticle reconstruction of the equatorial belt shows the outer genome spools (indexed -2 to 2) beneath the HBV capsid. The equatorial spool (indexed 0) lies under a hexameric patch of dimers (cyan hexagon), present at ten quasi-sixfold positions around the capsid. CTD density (red) projects from the inner capsid toward the outer spool, forming regularly spaced contact points along adjacent spools (red arrowheads). A side view of the hexameric patch (right) resolves CTD densities from the AB and CD dimers in contact with the spool DNA. Cyan hexagons and blue pentagons mark quasi-sixfold and fivefold positions, respectively. **b**, Top view shows a pentameric arrangement of CTD densities converging on the fivefold axis at the polar region of the viral particle. In the side view, CTD projections from AB dimers contact the outer spool through the central fivefold pore, and some CTD density appears smeared into the neighboring spool DNA (red arrowheads). Dashed orange brackets denote the spools indexed 5 and 6. At the fivefold, CTD density is radially concentrated toward the axis, approaching the center of the genome spool, whereas around quasi-sixfold belts it is more laterally distributed. These contacts are weak and are most apparent when the maps are rendered at lower contour levels. Together, these observations indicate that the CTD tethers and organizes the outer genome layer, with more continuous contacts around quasi-sixfold belts and sparser interactions at the fivefold pole, consistent with the increased mobility of the polar spool seen in Fig. 2.

An analogous situation has been seen with the dsRNA virus phi6. Phi6 also lacks a portal and has a coaxially spooled genome believed to be organized by interactions with positively charged capsid residues found in close proximity to the RNA.^17^ Another parallel of interest is with HIV, where reverse transcription of the genome is correlated with uncoating^58^ and, like with HBV, may act as a spring to destabilize the capsid^59^.

In summary, we have observed spooled dsDNA genomes in two non-portal viruses. Likely this is a shared feature among all hepadnaviruses. The spool is coaxial and may be oriented by capsid-DNA interactions. The exact basis of the spool’s fivefold orientation is unknown, but we speculate that the arrangement of the spool and its inherent energy plays a role in maturation signaling, capsid rupture, and the release of the viral genome from the capsid during infection.

## Methods

### Purification of Intracellular DHBV nucleocapsids

#### Cell culture

L59-3 is an LMH derivative which conditionally expresses the DHBV3 strain under control of TetRKrab fusion transcriptional repressor, such that expression of pgRNA is dependent on the presence of doxycyline. Cells were maintained in DMEM F12 (Gibco, Cat. 11-320-082) supplemented with 5% FBS (v/v) (Life Technologies, 16140-07) and 100 I.U./ml penicillin-streptomycin (Gibco, 15140-122). Flasks for passaging were coated with 0.1% (v/v) gelatin (Sigma-Aldrich G1393). For DHBV production, 14 T-175 flasks of L59-3 were grown in DMEM F12 with 5% FBS for 1 day and then DHBV production was induced for 8 days by adding 20 ng/ml Doxycycline to the media every other day. Flasks were washed twice with PBS (Gibco 10010-023) and stored at -80°C for later purification.

#### DHBV capsid purification via ultracentrifugation

Frozen L59-3 flasks were thawed and the cell lysates were harvested by incubation for 10 minutes at 37 °C with Core lysis buffer (50 mM Tris-HCl pH 8.0, 1% NP-40, 1 mM EDTA, 1× Pierce protease inhibitor, 10 mM NaF, 50mM β-glycerophosphate, 10 mM Na pyrophosphate, and DNase/RNase-free ultrapure water), 7 ml per flask. The lysate was centrifuged 3× at 10,000 × g at 4 °C for 25 minutes, transferring to a new conical tube between rounds. Supernatants were either treated after the first and second round of centrifugation with 20 µg/ml RNaseA or treated later after the linear gradient with 50 µg/ml RNaseA and 20 mM EDTA. RNaseA treatment was followed by incubation at 37 °C for 1 hour.

For capsid purification, the samples were loaded (15 ml sample/tube) on top of 5 ml 30% (w/v) sucrose cushion in SW32 tubes. All sucrose solutions for DHBV purification were prepared in buffer containing 100 mM Tris-HCL pH 8.0, 500 mM NaCl, 50 mM EDTA, 0.01% Triton X-100, 0.1% NP-40, 1× Pierce protease inhibitor, 10 mM NaF, 50mM beta-glycerophosphate, 10 mM Na pyrophosphate, and ultrapure water. Cushions were centrifuged at 110,880 × g for 8 hrs. The pellets were resuspended by overnight incubation, with stirring, in 650 µl buffer, homogenized on ice, treated with NP-40 (1% v/v), and sonicated in a water bath (15× at 10 seconds on, 10 seconds off). The sample was then centrifuged at 12,000 × g for 20 minutes at 4 °C

The supernatant was layered onto two linear sucrose gradients prepared in SW40 tubes by depositing 5 ml 40% (w/v) sucrose under 5 ml 5% (w/v) sucrose and spinning in a Biocomp gradient maker set to 5-40% short sucrose. The gradient was centrifuged at 92,262 × g for 4 hours at 4 °C using an SW40 Ti rotor. Fractions were collected in 1 ml increments from a hole pierced in the bottom of the tube and evaluated for DHBV DNA content by qPCR and for capsids by negative stain Transmission Electron Microscopy (TEM). Fractions 4-6 from each gradient were pooled together and diluted to 60 ml with buffer. The sample was then loaded over two 30% sucrose cushions in SW32 tubes and ran as before. The pellets were resuspended in 250 µl buffer overnight with stirring and then homogenized. The homogenizer was rinsed with 100 µl buffer, bringing the total volume of the combined sample to 600 µl. This was sonicated in a water bath (10× at 10 seconds on, 1 minute off) and clarified at 12,000 × g and 4 °C for 20 minutes. The sample was concentrated by loading on a 1 ml 30% sucrose cushion and spinning at 245,000 × g for 2.5 hrs at 4 °C in a TLS-55 rotor. The pellet was resuspended in 50 µl resuspension buffer (50mM Tris-HCl, 150 mM NaCl, and 2 mM DTT) and the ultracentrifuge tube was rinsed with an additional 20 µl resuspension buffer. The tube was then washed again with 70 µl resuspension buffer, keeping this wash sample separate from the pellet resuspension. After evaluating the sample clarity with negative stain TEM, we selected the wash sample for cryo-EM analysis.

### Purification of Intracellular HBV Capsids

#### Cell culture and transient transfection

HepG2 cells were cultured in DMEM/F12 (Gibco, Cat. 11-320-082) supplemented with 10% FBS (v/v) and 100 I.U./ml penicillin-streptomycin. To transfect HepG2 cells, the HBV infectious DNA clone HBV-HBc-WT plasmid^60^ was diluted to 0.01 µg/µl in serum-free Opti-MEM™ (Gibco, Cat. 31985070). The transfection reagent, X-tremeGENE™ HP DNA (Roche, Cat. 06366546001), was added at a 3:1 ratio (reagent to DNA) to the diluted plasmid solution and incubated at room temperature for 15 minutes. The transfection mixture was then added dropwise to 10-cm culture plates containing the cells. After incubation at 37°C overnight, the medium was replaced with fresh DMEM/F12 supplemented with 10% FBS and 100 I.U./ml penicillin-streptomycin.\

Media changes were performed every 2 days after transfection. On day 5 post-transfection, HepG2 cells were harvested for purification. The medium was removed via suction, and cells were washed three times with 1× HBSS. Plates were then allowed to dry, wrapped in foil, and either stored at -80°C for later purification or processed immediately for intracellular capsid purification.

#### HBV capsid purification via ultracentrifugation

Intracellular HBV capsids were purified from HepG2 cells using multiple rounds of ultracentrifugation. Frozen HepG2 cell plates were retrieved from -80°C storage and thawed to room temperature. Core lysis buffer (50 mM Tris, pH 8.0, 1% NP-40, 1 mM EDTA, 1× protease inhibitor, 10 mM NaF, 50 mM β-glycerophosphate, 10 mM Na pyrophosphate, 2 mM Na_3_VO_4_, and DNase/RNase-free ultrapure water) was added directly to the plates, followed by incubation at 37°C for 5 to 10 minutes. Cells were then scraped from the plates and transferred to a 50 mL ultra-high performance conical tube, and residual lysate was collected by washing the plates once more with core lysis buffer. The lysate was centrifuged at 11,635 × g for 20 minutes at 4°C, and the supernatant was transferred to a new conical tube.

To digest RNA, 200 μg/ml RNase A was added to the lysate and incubated at 37°C for 1 hour on a rotator. After incubation, the sample was centrifuged at 12,000 × g for 20 minutes at 4°C, and the supernatant was transferred to a new conical tube. To ensure complete RNA digestion, an additional round of RNase A treatment (0.1 mg/µl) was performed, followed by incubation at 37°C for 1 hour and centrifugation at 12,000 × g for 20 minutes at 4°C.

For capsid purification, a 10%/20% (w/v) step sucrose gradient was prepared in an open-top ultra-clear tube, and the sample was carefully layered onto the gradient by pipetting. The gradient was ultracentrifuged at 124,921 × g for 15 hours at 4°C using a Beckman Coulter ultracentrifuge with SW32-Ti buckets. The resulting pellet was resuspended and homogenized on ice until fully dissolved. To remove residual nucleic acids, 5 mM MgOAc and 200 µg/ml DNase I were added to the homogenized sample, followed by incubation at 37°C for 30 minutes. Subsequently, 20 mM EDTA and 200 µg/ml RNase A were added, and the mixture was incubated at 37°C for 1 hour. The reaction was then centrifuged at 12,000 × g for 20 minutes at 4°C.

The supernatant was layered onto a 15–30% (w/v) continuous sucrose gradient containing 20 mM Tris-HCl (pH 7.5), 50 mM NaCl, 1 mM EDTA, 0.01% (v/v) Triton X-100, 0.1% (v/v) NP-40, 2.5× protease inhibitor, 0.1 M NaF, 0.1 M β-glycerophosphate, 10 mM Na pyrophosphate, 5 mM Na_3_VO_4_, 0.05% (v/v) β-mercaptoethanol, and ultrapure water. The gradient was ultracentrifuged at 124,921 × g for 4 hours at 4°C using an SW32 Ti rotor, and the resulting fractions were collected on ice for further analysis. Fractions containing viral DNA signals were pooled and concentrated using Nanosep for the subsequent EM and cryo-EM imaging.

### Identification of HBV Capsid Protein / pgRNA / DNA

#### Native agarose gel electrophoresis (NAGE)

A NAGE was performed to assess the presence of capsids in the sucrose gradient fractions. In brief, a 1% (w/v) agarose gel was prepared by dissolving agarose powder in 1× TAE (Tris-acetate-EDTA, Invitrogen^™^) and allowing it to solidify. The loading samples were prepared by mixing each fraction with 6x SDS-free loading dye, which contains 30% (v/v) glycerol, 0.25% (w/v) bromophenol blue, 0.25% (w/v) xylene cyanole FF and ultrapure distilled water (dH_2_O). The samples were then loaded into the wells, and electrophoresis was carried out at 100-130V for 3-4hr.

#### Immunoblot analysis

HBV capsids in sucrose gradient fractions were analyzed by western blot. Following SDS-PAGE, proteins were transferred onto a 0.45 µm nitrocellulose membrane using filter paper support. The membrane was incubated overnight at 4 °C with a primary anti-HBcAg antibody (Zeta, Cat. Z2085RL-A), followed by incubation with an HRP-conjugated goat anti-rabbit secondary antibody (Invitrogen, Cat. P-2271MP) for 1 h at room temperature on a rocker. After three washes, signals were developed and imaged using a ChemiDoc MP system (Bio-Rad) with Image Lab software. Custom 32P-labeled pgRNA- and HBV DNA-specific probes were used to detect RNA and DNA fractions, respectively.

### Negative-stain Transmission electron microscopy (TEM)

TEM grids were prepared to assess capsid concentration and quality. Carbon-coated grids (EMS, Cat. CF300-Cu-50) were glow-discharged carbon-side up to improve sample adhesion. A 4 µl of sample solution was applied to the grids and incubated for 30 seconds, followed by blotting with filter paper to remove excess liquid. To remove residual sucrose buffer, 4 µl of dH_2_O was added to the grids and immediately blotted with filter paper; this step was repeated twice for HBV samples.

For staining, 4 µl of 2% (w/v) uranyl acetate was applied to the grids and incubated for 20-30 seconds, then blotted with filter paper to remove excess stain. The grids were allowed to air dry and were subsequently imaged using a JEOL-1400 TEM operated at 60 kV equipped with an AMT NanoSprint43M-Mk2 CMOS camera (for HBV samples) or a JEOL-1400plus TEM operated at 120 kV equipped with a Gatan OneView CMOS camera (for DHBV samples).

### Preparation of cryo-EM samples and imaging

HBV and DHBV grids were prepared by glow discharging EMS ultrathin continuous, 300 mesh, carbon film coated grids, and depositing 4 µl sample prior to vitrification using a Thermo Fisher Scientific Vitrobot (Mark IV). 16,406 (DHBV) and 17,424 (HBV) movies were collected on a Titan Krios electron microscope at 300 kV equipped with a Gaton K3 camera in Super Resolution Counting Mode. Movies were collected at 64,000 × magnification with an exposure of 30 e−/Å^2^ (∼ 1 e−/frame). Defocuses values were −6.2 µm to −0.3 µm (DHBV) and −5.4 µm to −0.5 µm (HBV).

### Image processing

Image processing was performed in RELION. Movies were motion corrected and CTF estimation was performed with CTFFIND 4.1. Images with a max CTF resolution 6 Å and under were retained (leaving 15,991 DHBV movies and 16,071 HBV movies). After auto-picking particles, well-resolving particles were selected by 2D classification and subjected to 3D refinement, CTF refinement, particle polishing, and post processing to reach a final resolution of 3.3 Å (DHBV, 122,883 particles) and 3.5 Å (HBV, 18,257 particles) with icosahedral (I2) symmetry imposed.

### Asymmetric reconstruction of DHBV genome density

The workflow is illustrated in Extended Data Fig. 2. DHBV particles were sorted by 3D classification with I2 symmetry imposed. Each class (except the smallest class with 6619 particles) went through 3D refinement and was then expanded according to I2 symmetry using relion_particle_symmetry_expand. Each was then subjected to one or more rounds of 3D classification without imposed symmetry, using a mask to exclude capsid density. Classes with apparent DNA rings were selected for the final round of masked 3D classification or subjected to additional rounds of 3D classification first. The final selection—375,165 expanded particles—were aligned together using EMAN2 and starpy before classification into two classes. For the asymmetric structure of the spooled genome plus the capsid, the full map was generated from the better resolved 3D class using relion_reconstruction and then 3D refined with local refinement only (0.8/0.8 angular sampling). When used in figures, this map was lowpassed to two stranded deviations. The 3D class without any further processing was used for DHBV DNA in Fig. 1 and 3.

### Asymmetric reconstruction of HBV genome density

To resolve the nucleic acid density, we used a strategy that broke symmetry from the outside in. A mask that encloses the first layer of nucleic acid (Extend Data Fig. 4) was applied. Symmetry expansion and focused classification revealed spooling double-stranded DNA density at this layer. To resolve the conformation of the second layer, the particle orientations from previously observed spooling dsDNA density were kept fixed, and then focused classification was applied to tease out the conformational heterogeneity at the second DNA layer (gold color). Because the outer DNA conformation was fixed, we could clearly observe several double-stranded spooling densities at the second layer as well. One of the particle groups was selected and further refined asymmetrically to a final reconstruction at 4 Å. To resolve the inner core structure, an additional 3D classification was applied with a mask that enclosed the inner core. The final internal core showed partially double-stranded and partially single-stranded DNA density, with additional density at the center of the particles.

## Data Availability

The cryo-EM density maps has been deposited to the EM Data Bank with accessions numbers EMDB-73588 (DHBV capsid), EMD-73598 (C1 DHBV capsid with DNA), EMD-71261 (HBV capsid), and EMD-71262 (C1 HBV capsid with DNA). Protein models have been deposited to the Protein Data Bank under accession numbers 9YX3 (DHBV capsid) and 9P42 (HBV capsid).

## Acknowledgements

This research was funded by National Institute of Allergy and Infectious Diseases NIH grants R01AI173104 (J.W.) R01AI144022 (A.Z), R37AI043453 (J.H), and R01AI060018 (D.L), as well as National Cancer Institute NIH grant P01CA022443 (D.L.).

## Contributions

J.W. and A.Z. conceived and jointly supervised the study. N.G. contributed to development of analyses, performed experiments and data analysis for DHBV, and generated figures and tables for the manuscript.

J.W. performed experiments and data analysis and generated figures for the manuscript. KC worked on data analysis. H.L. contributed to HBV expression and purification, and J.X. assisted with HBV expression and purification. S.N. contributed to developing DHBV expression, D.L. and K.P. provided support and the L59 cell line. JH provided cell lines and advice for HBV expression and purification. N.G., A.Z., and J.W. wrote and revised the manuscript, and all co-authors reviewed the manuscript.

---

**Extended Data Fig. 1.**
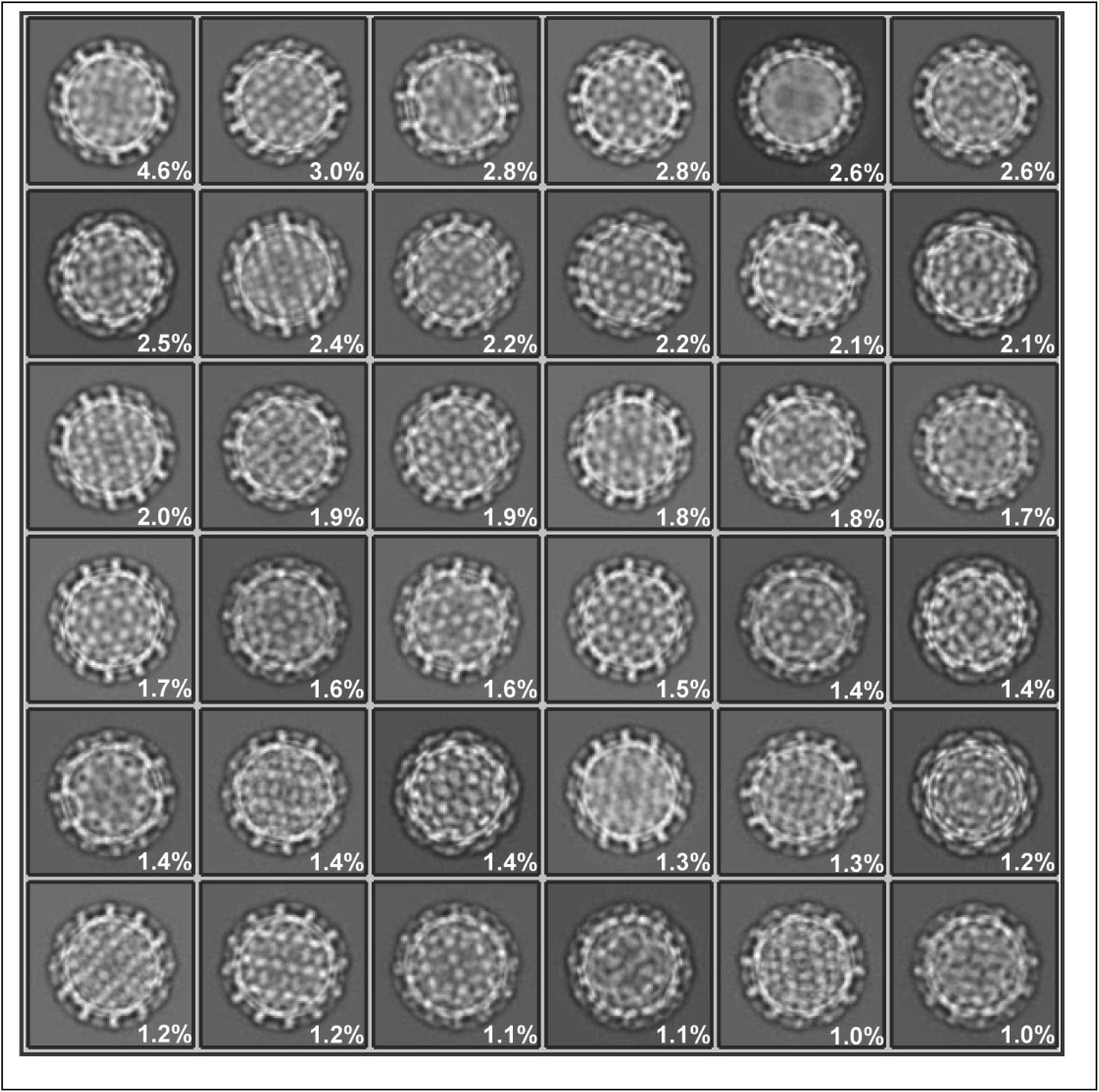
The top 36 2D classes used for reconstruction of the DHBV capsid. Class occupancy out of 122,883 particles is shown by percentage.

**Extended Data Fig. 2.**
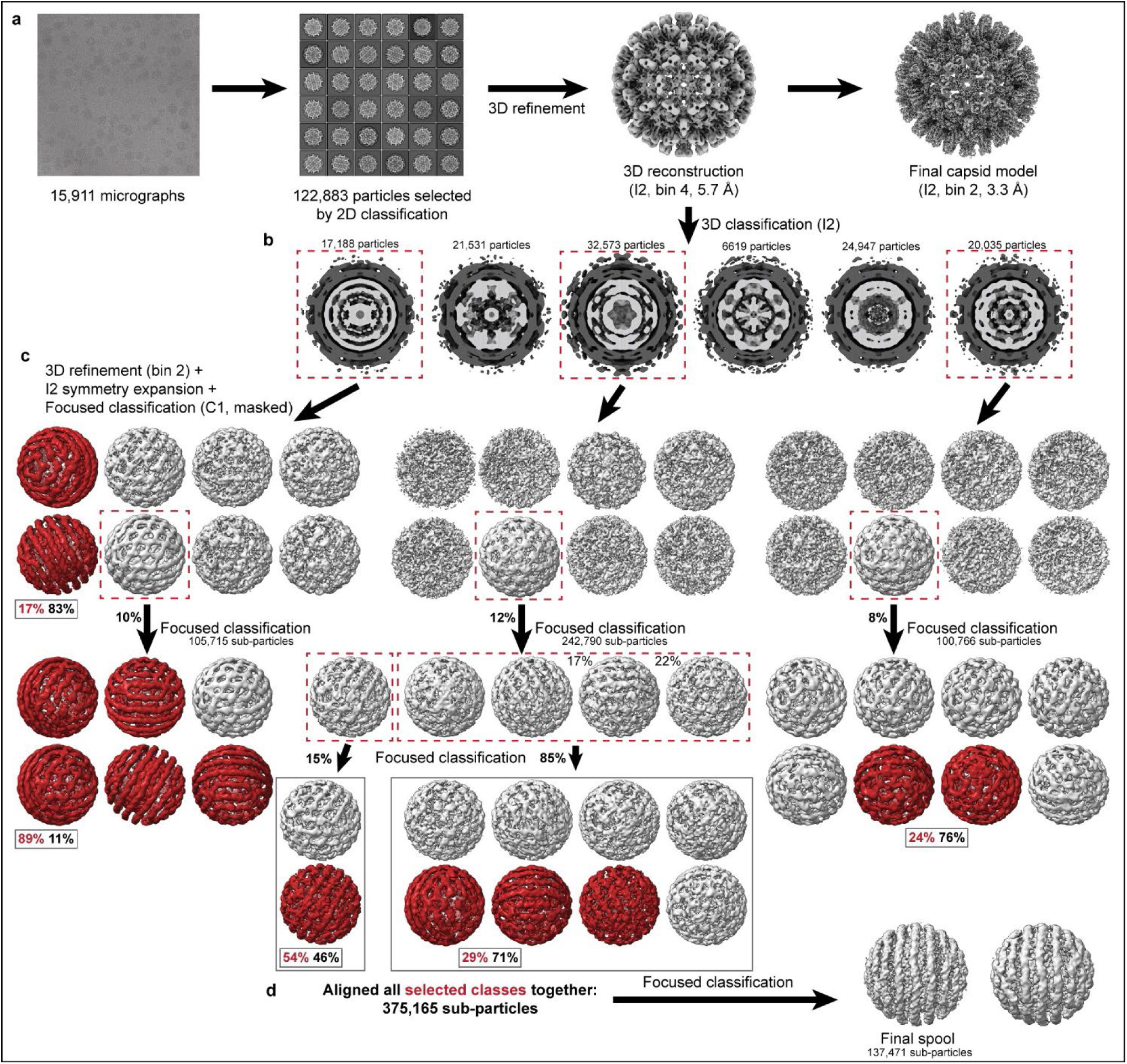
Data processing workflow for DHBV nucleocapsids and genome. **a**, Reconstruction of icosahedral (I2) capsid **b,** Classes resulting from whole particle 3D classification with I2 symmetry imposed. Capsid density is indicated in dark grey vs. internal/genomic density (light grey) **c**, Sets of classes resulting from masked asymmetric classification with subparticles generated from classes selected from **(b)**. Classes indicated with dashed boxes were subjected to additional rounds of classification. Boxed percentages (red) indicate the percent of subparticles from that classification set used for the final structure in **(d)** With the exception of the smallest class with 6619 particles which we did not further classify, the other two classes in **(b)** for which further 3D classification is not shown did not yield spool-like classes. **d**, Final spool structure generated from masked asymmetric classification of particles from classes colored red in **(c)**

**Extended Data Fig. 3.**
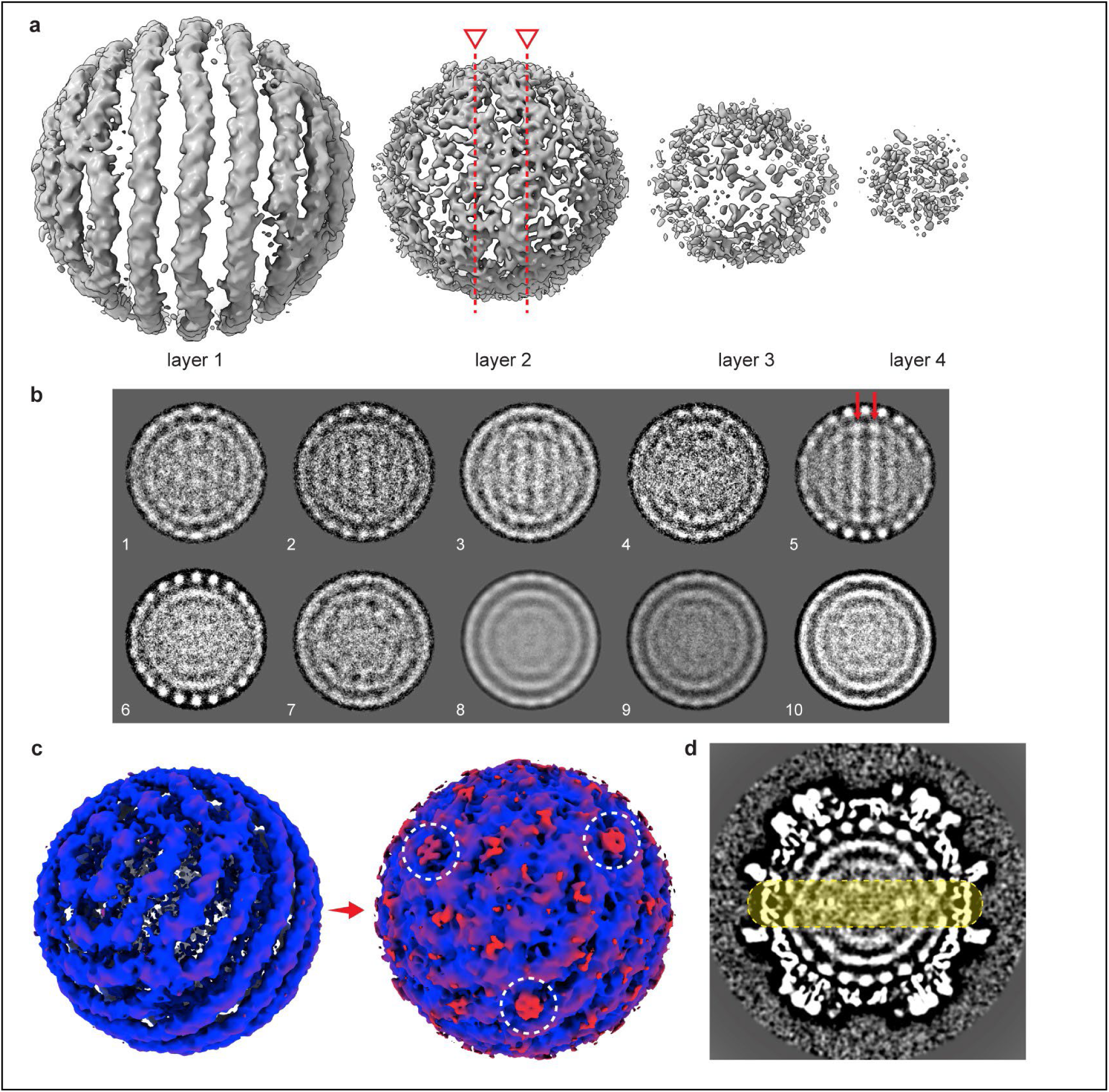
Layers and density connections in the DHBV spool reconstruction. **a**, Layers of density in DHBV spool structure. The back of the density is not shown. The layers include density > 106 Å from center (layer 1), 79-106 Å (layer 2), 48-79 Å (layer 3), and 0-48 Å from center (layer 4). Red lines indicate the position of the central two apparent DNA strands in the second layer. **b**, 2D classification of spool density after signal subtraction. Red arrows in class 5 indicate suggested second layer strands, also indicated in **(a)**. **c**, Spool structure colored by radial position to emphasize positions where density protrudes from the structure (red, circled in white). Protrusions are visible only at low density contour (right) and are found at the 5-fold symmetry positions. **d**, Orthoplanes representation of the asymmetric reconstruction including both the capsid and spool. The position of the density connections between layers underneath the capsid 5-folds (plus the protrusions from the spool) are indicated with a yellow circle.

**Extended Data Fig. 4.**
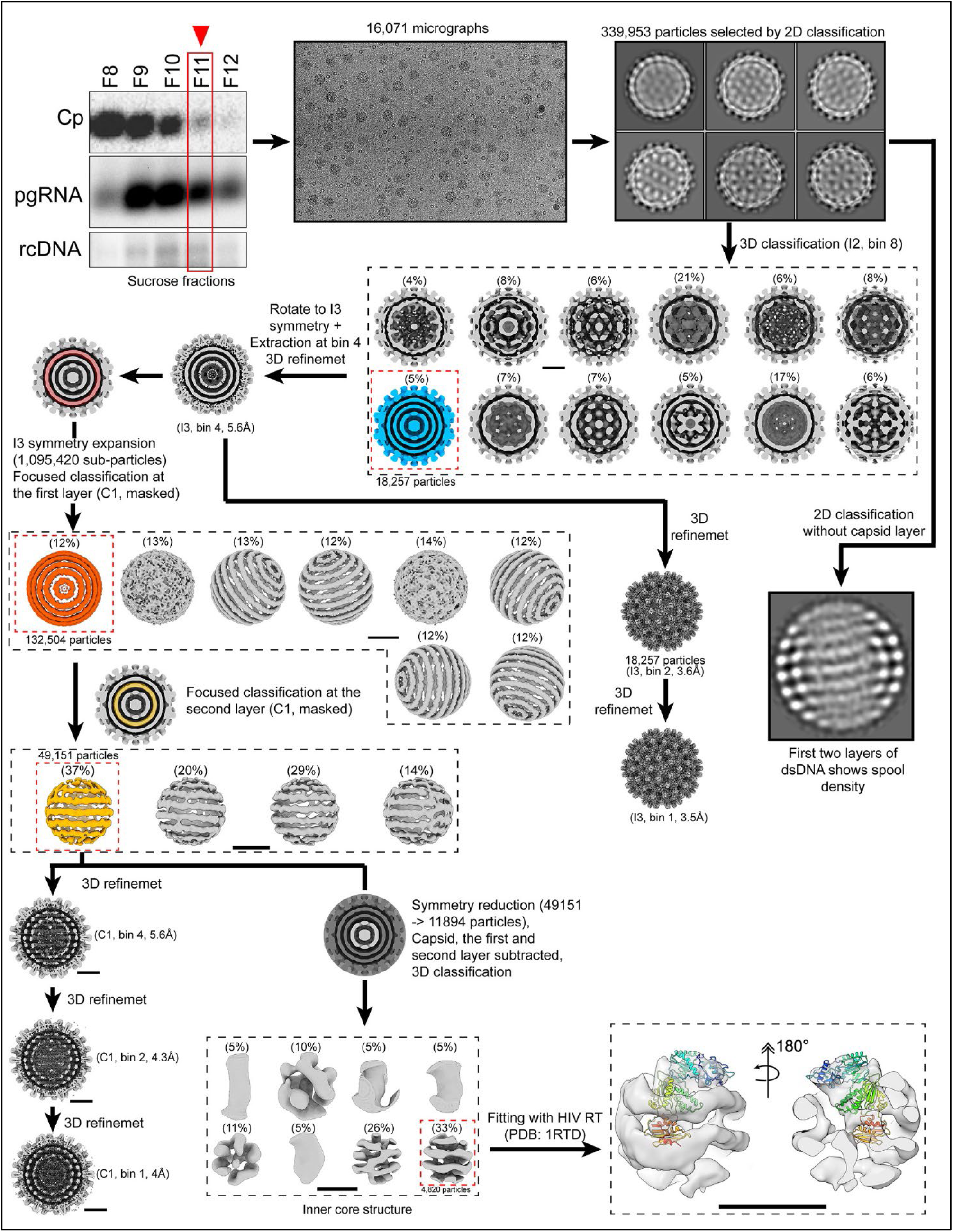
Data processing workflow for rcDNA-filled HBV nucleocapsids and genome layers. A workflow illustrates the cryo-EM data processing pipeline for capsid and genome reconstruction. The sample fraction containing intracellular HBV capsids, pgRNA, and rcDNA was selected for cryo-EM data collection. Images were motion-corrected, CTF-estimated, and subjected to iterative 2D classification (bin 8, data compression level) to remove contaminants. An *ab initio* 3D model served as a reference for 3D classification to separate empty, immature, and mature particles. Classes outlined with dashed black lines indicate 3D classification results; the class enclosed by red dashed lines was selected for further refinement. The class resembling duck rcDNA-filled capsids, showing concentric nucleic acid rings, was refined under icosahedral (I3) symmetry to 5.6 Å (bin 4). Subsequent processing was divided into two paths. One continued icosahedral refinement, yielding a 3.5 Å map. The second used an asymmetric, layer-by-layer strategy to dissect the rcDNA structure. Focused masking on the first nucleic acid layer (dark orange), followed by symmetry expansion and classification, revealed spooled dsDNA density. Fixing the first-layer orientations enabled focused classification of the second layer (gold), revealing distinct dsDNA spooling patterns. A separate 2D analysis was also performed. Upon radial removal of the capsid density, 2D classification of the isolated genome particles revealed two spooled DNA layers, in agreement with the 3D reconstructions. One particle group was further refined asymmetrically to 4 Å, while additional 3D classification resolved the internal core containing partially double- and single-stranded DNA with extra central density. This central feature can accommodate a reverse transcriptase–like structure modeled from HIV RT (PDB 1RTD).

**Extended Data Fig. 5.**
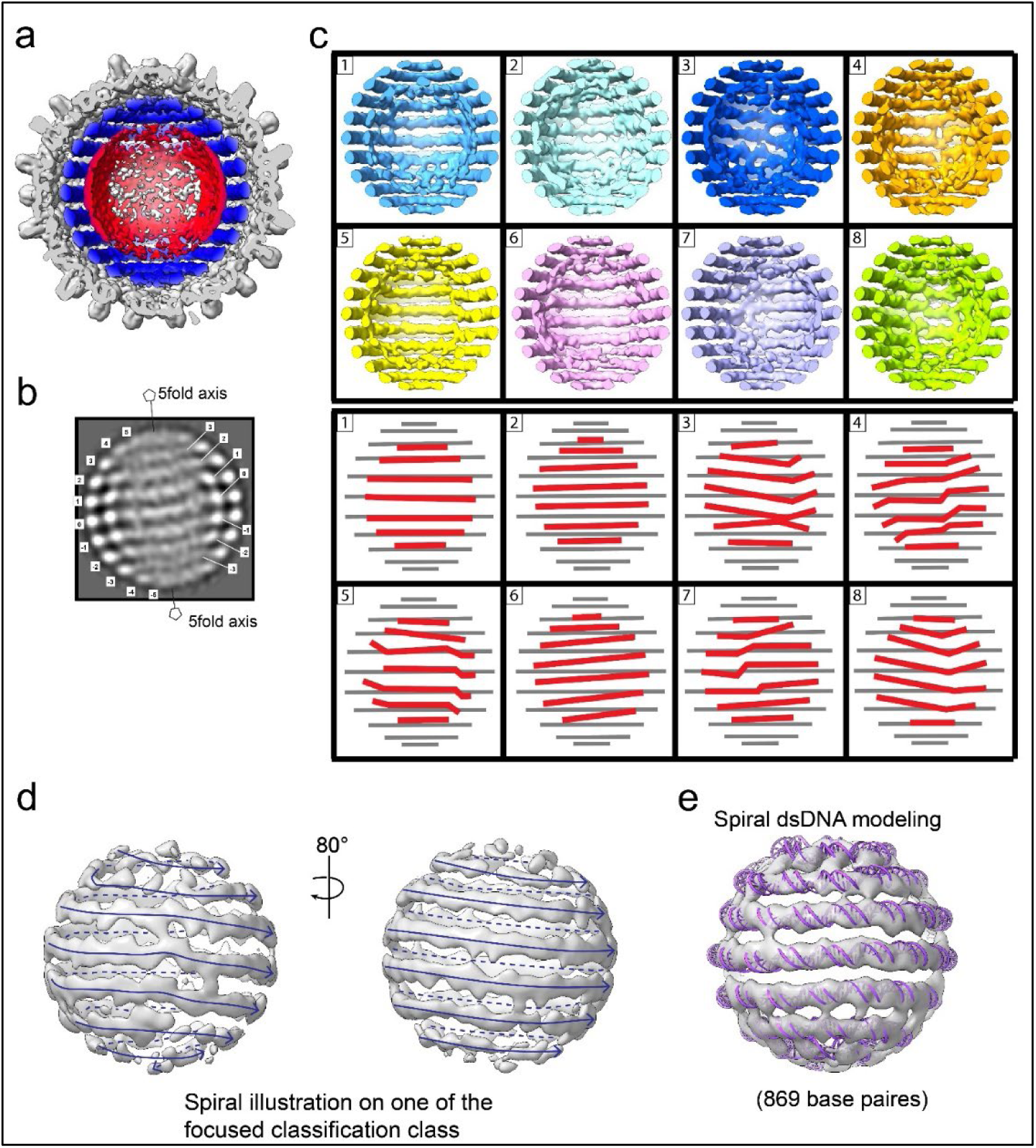
Structural analysis of the second layer of HBV genome. (**a**) When the first DNA layer (blue) is aligned relative to the icosahedral capsid, density from the second layer and inner core (red) averages into a smear, indicating flexibility relative to the outer layer. (**b**) To validate internal spool-like genome density, the capsid signal of each particle was computationally masked, and 2D classification was performed. The resulting class average clearly shows spool-like rings at both the first and second layers inside the capsid, with the number of rings matching the 3D reconstructions. The second layer lies beneath the first but displays non-uniform curvature; weak densities appear to connect layers along the polar axis. (**c**) To assess this flexibility, orientations of the outer layer were first fixed, followed by 3D classification without image alignment on the second layer. The resulting classes (upper panels) place the second layer beneath or between strands of the outer layer; cartoons (lower panels) illustrate the outer DNA in grey and the second layer in red. Several classes exhibit continuous, spiral paths rather than discrete ring-like spools. (**d**) Representative focused classified second-layer cryo-EM map rendered as a continuous spiral (two views, separated by 80°). (**e**) Ideal B-form dsDNA fitted to the continuous-spiral density (∼869 base pairs), consistent with the observed periodicity and groove register.

**Extended Data Fig. 6.**
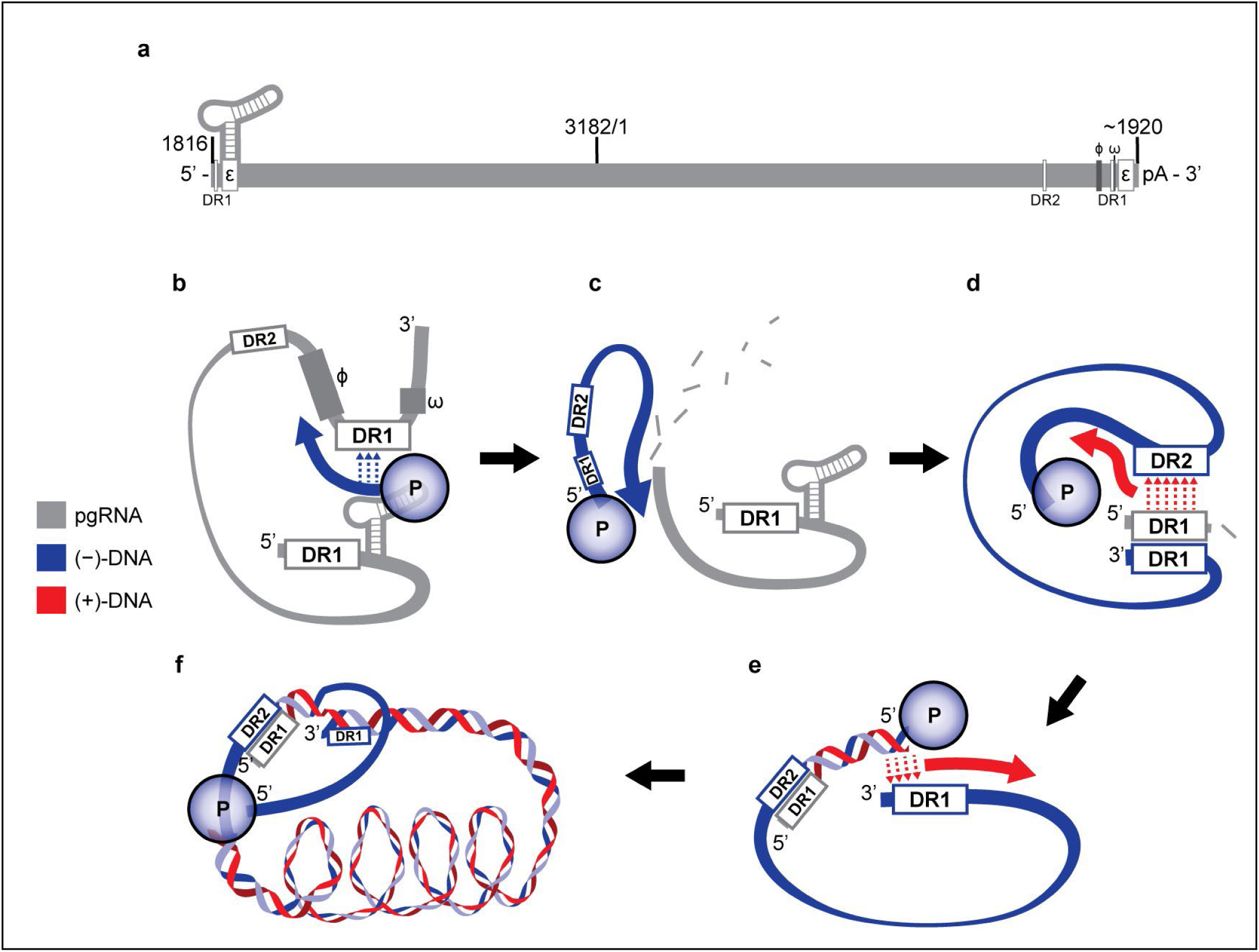
Reverse transcription in HBV. **a**, pgRNA transcript, approximately to scale, numbered with the EcoRI cutsite at position 3182/1. DR1 and DR2 refer to direct repeat sequences, and ϕ and ω are cis-acting sequences involved in directing DNA synthesis. The genome is terminally redundant. **b**, Reverse transcription of (−)-DNA (blue) begins with the synthesis of a three-nucleotide primer using ε as a template. The first template switch consists of this primer aligning with a complementary sequence within DR1, allowing (−) strand synthesis to continue, with polymerase (P) remaining covalently bound to the (−) strand^61^. Base pairing between ϕ, ω, and ε may help facilitate the switch^62^. **c**, The P RNaseH domain degrades pgRNA during (+) strand synthesis. **d**, In the second template switch (primer translocation), DR1 at the 5’ end of pgRNA complements with the newly synthesized DR2 on the (−) strand, allowing P to synthesize (+)-DNA using the (−) strand as a template. **e**, In the third template switch, the repeat region at the 3’ end of the (+)-DNA complements with DR1 at the 3’ end of (−)-DNA, allowing continued (+) strand synthesis and circularization of the genome. **f**, (+)-DNA synthesis continues towards the 5’ end of the (−) strand template, but does not complete, leaving a partially single stranded circular genome.

**Extended Data Fig. 7.**
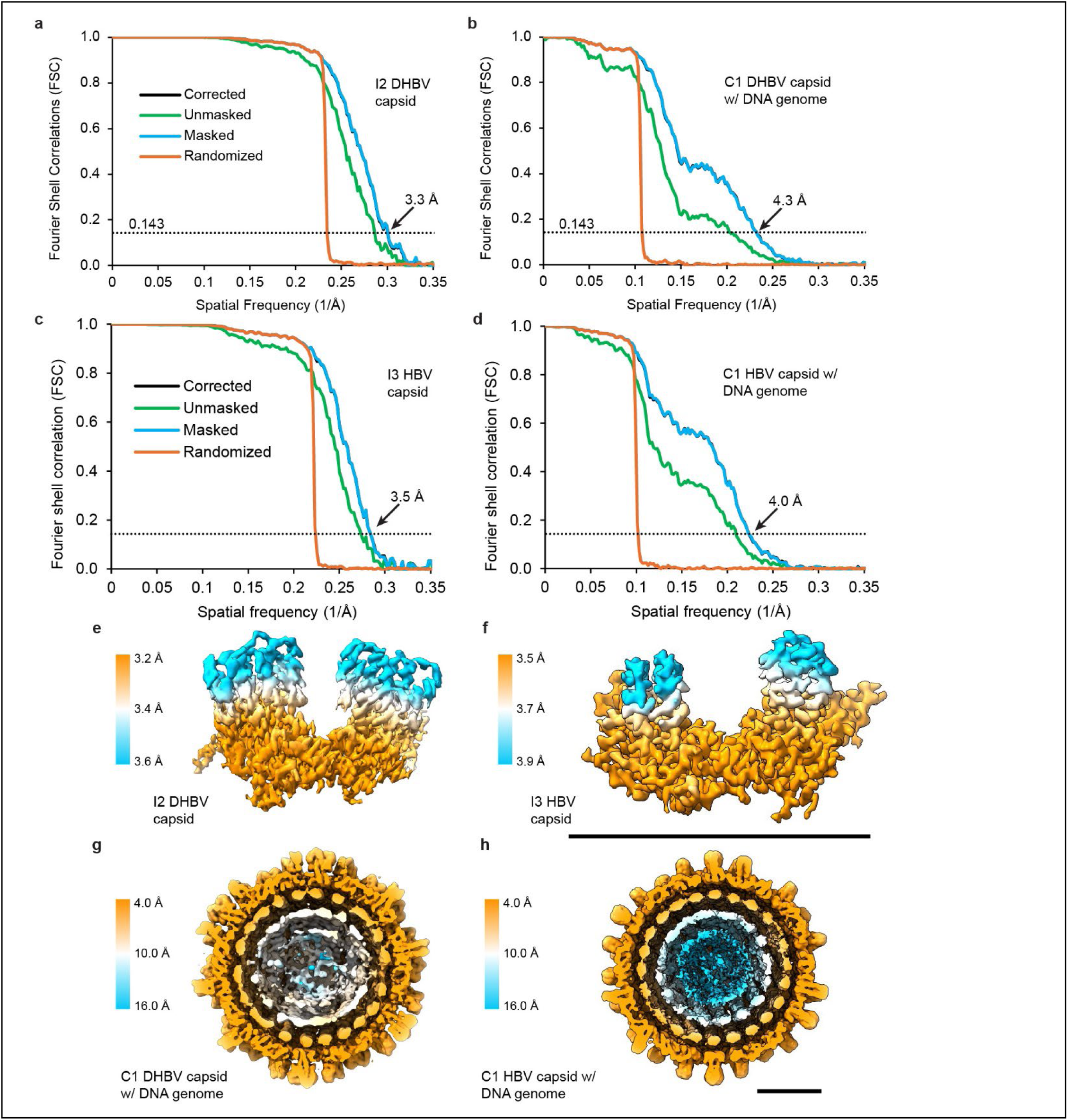
HBV and DHBV structure resolution. **a-d**, Gold-standard Fourier Shell Correlation (FSC) of DHBV and HBV reconstructions **e**,**f**, DHBV and HBV subunit unit density from the icosahedral (I2 and I3) capsid reconstructions, colored by local resolution calculated in RELION. **g**,**h**, Density cross-section of the DHBV and HBV asymmetric capsid with DNA genome reconstructions, colored by local resolution calculated in Relion

**Extended Data Table 1.**
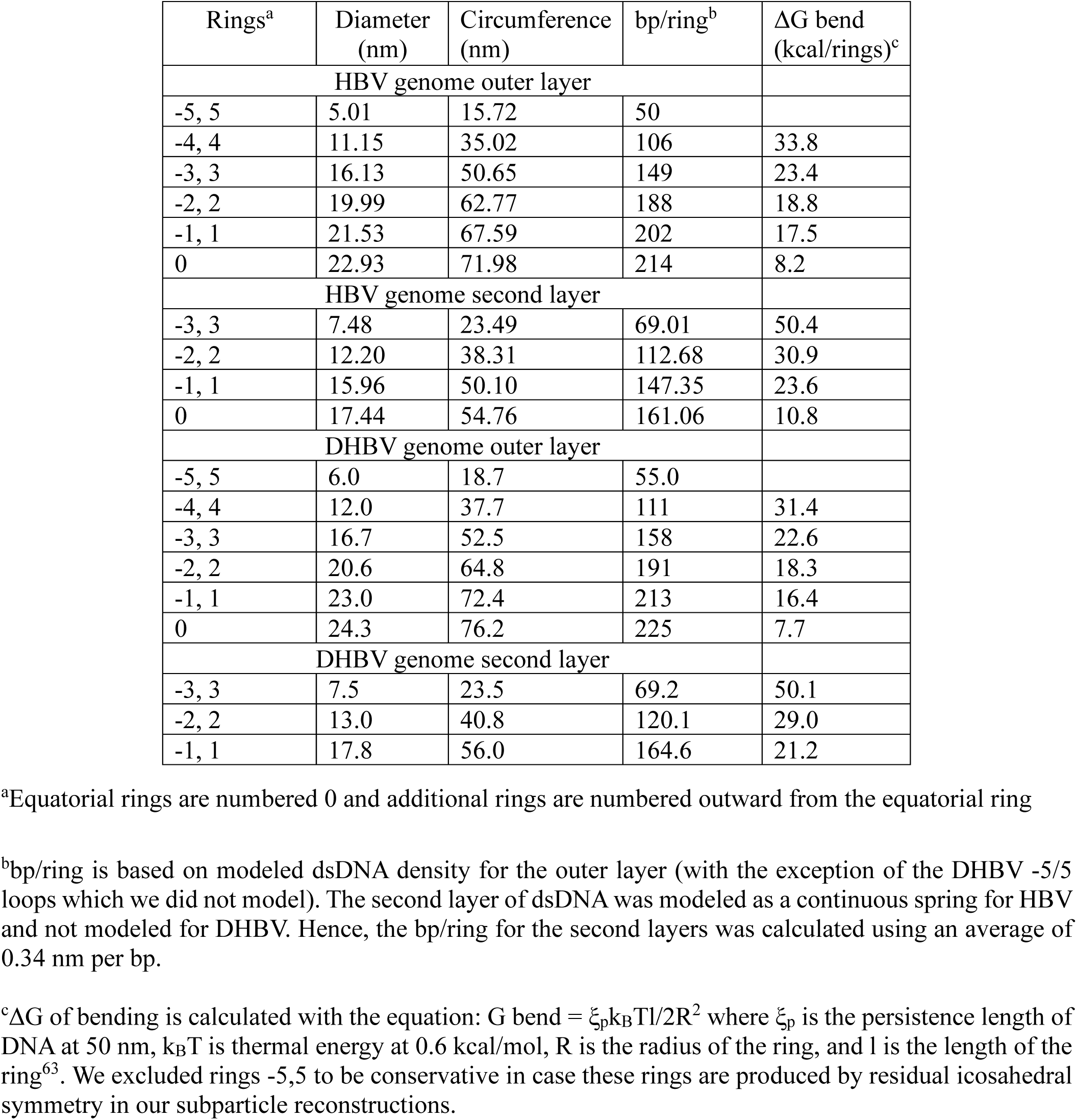
DNA accommodated by first and second HBV and DHBV genome layers.

| Rings<sup>a</sup> | Diameter (nm) | Circumference (nm) | bp/ring<sup>b</sup> | ΔG bend (kcal/rings)<sup>c</sup> |
| --- | --- | --- | --- | --- |
| HBV genome outer layer |  |  |  |  |
| -5, 5 | 5.01 | 15.72 | 50 |  |
| -4, 4 | 11.15 | 35.02 | 106 | 33.8 |
| -3, 3 | 16.13 | 50.65 | 149 | 23.4 |
| -2, 2 | 19.99 | 62.77 | 188 | 18.8 |
| -1, 1 | 21.53 | 67.59 | 202 | 17.5 |
| 0 | 22.93 | 71.98 | 214 | 8.2 |
| HBV genome second layer |  |  |  |  |
| -3, 3 | 7.48 | 23.49 | 69.01 | 50.4 |
| -2, 2 | 12.20 | 38.31 | 112.68 | 30.9 |
| -1, 1 | 15.96 | 50.10 | 147.35 | 23.6 |
| 0 | 17.44 | 54.76 | 161.06 | 10.8 |
| DHBV genome outer layer |  |  |  |  |
| -5, 5 | 6.0 | 18.7 | 55.0 |  |
| -4, 4 | 12.0 | 37.7 | 111 | 31.4 |
| -3, 3 | 16.7 | 52.5 | 158 | 22.6 |
| -2, 2 | 20.6 | 64.8 | 191 | 18.3 |
| -1, 1 | 23.0 | 72.4 | 213 | 16.4 |
| 0 | 24.3 | 76.2 | 225 | 7.7 |
| DHBV genome second layer |  |  |  |  |
| -3, 3 | 7.5 | 23.5 | 69.2 | 50.1 |
| -2, 2 | 13.0 | 40.8 | 120.1 | 29.0 |
| -1, 1 | 17.8 | 56.0 | 164.6 | 21.2 |

<sup>a</sup>Equatorial rings are numbered 0 and additional rings are numbered outward from the equatorial ring
<sup>b</sup>bp/ring is based on modeled dsDNA density for the outer layer (with the exception of the DHBV -5/5 loops which we did not model). The second layer of dsDNA was modeled as a continuous spring for HBV and not modeled for DHBV. Hence, the bp/ring for the second layers was calculated using an average of 0.34 nm per bp.
<sup>c</sup>ΔG of bending is calculated with the equation: $G \text{ bend} = \xi_p k_B T l / 2R^2$ where $\xi_p$ is the persistence length of DNA at 50 nm, $k_B T$ is thermal energy at 0.6 kcal/mol, $R$ is the radius of the ring, and $l$ is the length of the ring<sup>63</sup>. We excluded rings -5,5 to be conservative in case these rings are produced by residual icosahedral symmetry in our subparticle reconstructions.

**Extended Data Table 2.** virus genome packing densities compared to RNA/DNA layer spacing and interhelix spacing.

| Virus | $\rho_{\text{pack}}$<br>(packing density)<sup>a</sup> | Lumen diameter<br>(nm)<sup>b</sup> | Volume<br>(nm<sup>3</sup>)<sup>c</sup> | Genome Length (kbp) | Layers Spacing (Å) | Strand Spacing (interhelix) ( Å) |
| --- | --- | --- | --- | --- | --- | --- |
| Phage Phi6 | 0.30 | 44.9 | 47,522 | 13.4 | 29<sup>d</sup> | 26<sup>d</sup> |
| DHBV | 0.34 | 26.2 | 9417 | 3 | 29-30<sup>e</sup> | 32<sup>e</sup> |
| HSV | 0.36 | 955 | 455,900 | 151 | 26-27<sup>f</sup> | 31-33<sup>f</sup> |
| HBV | 0.43 | 24.7 | 7919 | 3.2 | ~30<sup>e</sup> | 31<sup>e</sup> |
| Phage T4 | 0.52 | 87 | 345,600 | 169 | 23-25<sup>g</sup> | 26-29<sup>g</sup> |
| Phage T7 | 0.57 | 52 | 75190.87 | 39.9 | 22<sup>i</sup> | 26<sup>i</sup> |
| Phage T5 | 0.58 | 75 | 225,032.26 | 121.8 | 23<sup>i</sup> | 26-27<sup>i</sup> |
| Phage Phi29 | 0.65 | 39.2 | 31,544.59 | 19.3 | 23<sup>j</sup> | 25<sup>j</sup> |

<sup>a</sup> $\rho_{\text{pack}}$ is calculated by dividing virus lumen space (nm<sup>3</sup>) by genome length in nm ( $0.34\pi N_{\text{bp}}$ )<sup>63</sup>.
<sup>b</sup>For DNA phages, this is an approximate diameter based on the calculated volume.
<sup>c</sup>For Phages T5, T7, and Phi29, we used lumen volume estimates calculated by Bores and colleagues<sup>18</sup>. For HBV, DHBV, HSV (EMDB 9864), and the dsRNA phage Phi6 (EMDB 0302), lumen volumes were calculated by measuring the diameter of the packaged genome density. For Phage T4 (EMDB 32109), we approximated the capsid volume as a cylinder with two capping cones, and subtracting the intruding portal complex volume (approximated as a cylinder) from this.
<sup>d</sup>Data from Ilca et al.
<sup>e</sup>Measured from structures in this paper. Layer spacing is between the first and second genome layers and strand spacing is between rings in the outer layer.
<sup>f</sup>Measured from EMDB 9864. We measured strand spacing as between rings in the outer layer.
<sup>g</sup>Layer spacing in prolate heads was reported as 25 Å, corresponding to 28.9 Å with hexagonal packing<sup>47</sup> (for hexagonal packing, interhelix spacing = layer spacing $\times \frac{2}{\sqrt{3}}$ )<sup>43</sup> However, ~23 Å spacing is also reported between layers in isometric T4 capsids<sup>46</sup> (23.6) and in prolate diffraction data (22.9),<sup>64</sup> which would correspond to 26.4 hexagonal inter helix spacing.
<sup>h</sup>The reported 25.66 interhelix spacing for Phage T7<sup>65</sup> would correspond to 22.2 layer spacing with hexagonal packing.
<sup>i</sup>Hexagonal interhelix spacing for Phate T5 is reported as 26.4 Å<sup>20</sup> and 26.31-26.71 Å<sup>65</sup>, which corresponds to 22.79-23.13 layer spacing.
<sup>j</sup>Data from Bores et al.<sup>18</sup> and Woodson et al.<sup>66</sup>

**Extended Data Table 3.**
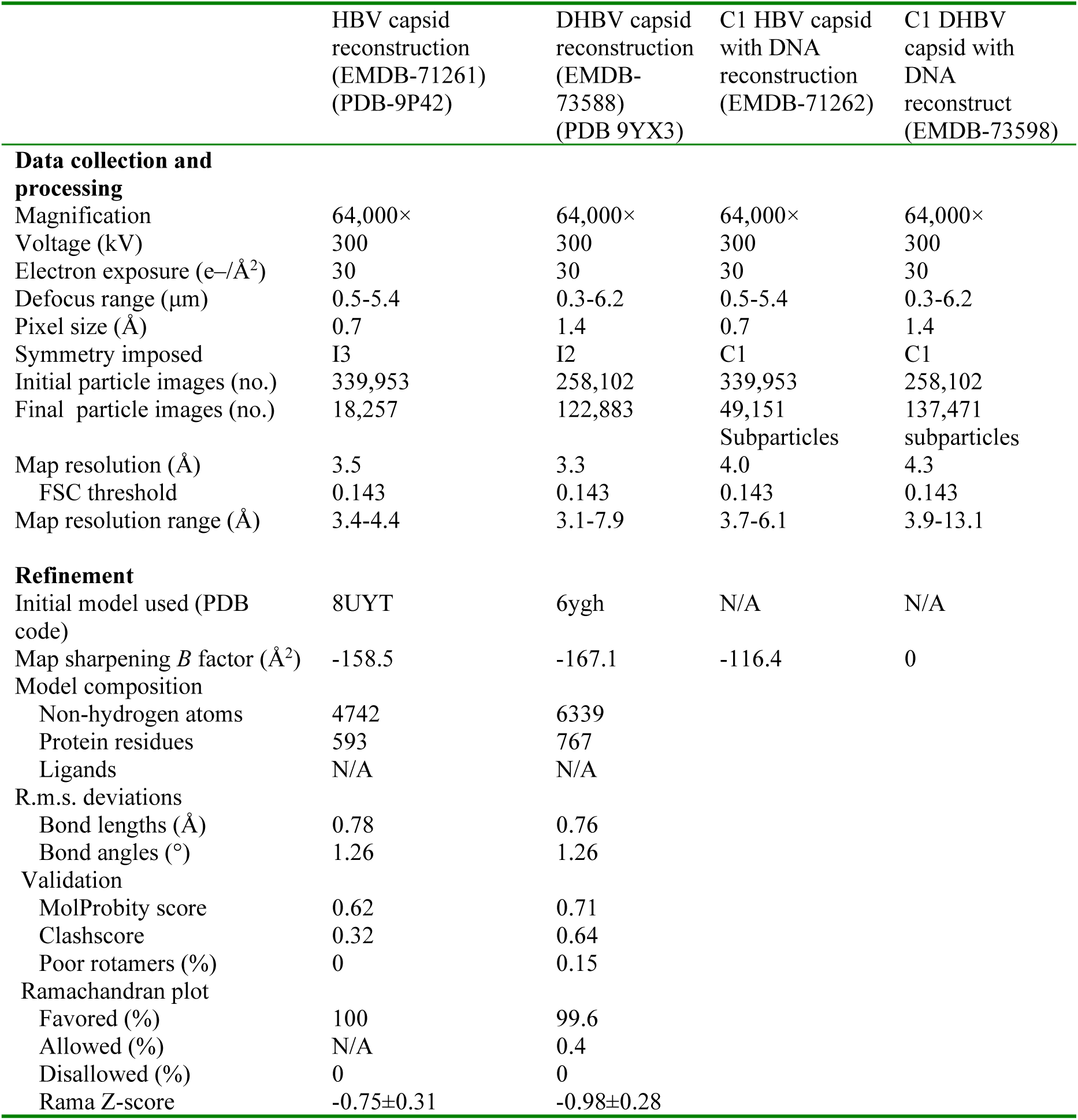
Cryo-EM data collection, refinement and validation statistics.

